# Spreading of molecular mechanical perturbations on linear filaments

**DOI:** 10.1101/573261

**Authors:** Zsombor Balassy, Anne-Marie Lauzon, Lennart Hilbert

**Affiliations:** Department of Physiology, McGill University, H3G 1Y6 Montreal, QC, Canada; Department of Biomedical Engineering, McGill University, H3A 2B4 Montreal, QC, Canada; Meakins-Christie Laboratories, Research Institute of the McGill University Health Centre, QC H4A 3J1 Montreal, QC, Canada; Department of Medicine, McGill University, H4A 3J1 Montreal, QC, Canada; Institute of Toxicology and Genetics, Karlsruhe Institute of Technology, 76021 Karlsruhe, Germany; Zoological Institute, Systems Biology and Bioinformatics, Karlsruhe Institute of Technology, 76131 Karlsruhe, Germany; Center for Systems Biology, 01307 Dresden, Germany; Max Planck Institute for the Physics of Complex Systems, 01187 Dresden, Germany; Max Planck Institute of Molecular Cell Biology and Genetics, 01307 Dresden, Germany

## Abstract

Global changes in the state of spatially distributed systems can often be traced back to events resulting from local interactions. Whether the results of local interactions grow into global changes, however, depends (i) on the system geometry and (ii) the spatial spreading of the outcomes of local interactions. Here, we investigate how different spreading behaviors of local events determine their global impact in one-dimensional systems of different size. In particular, we combine *in vitro* experiments where groups of myosin motors propel actin filaments, single-molecule resolution simulations of these *in vitro* experiments, and an abstracted spin chain model. All three approaches lead to the same two conclusions. First, local events that become long-term stable only after they have spread to full system size have more impact in smaller systems. Second, local events that are relatively stable upon initial occurrence and then spread to full system size have more impact in larger systems. Our work provides highly specific predictions for future experiments that resolve actin-myosin-crosslinker interactions along actin filaments. Also, the conclusions from our work should generally apply to local-to-global spreading in finite, one-dimensional geometries.

**Significance Statement:** We address the fundamental question of how results of local interactions spread in one-dimensional systems of different size. To this end, we reconstituted the molecular contractile machinery of muscle, which is organized around linear actin filaments of different length and drives their forward sliding. In addition, we use detailed simulations that follow the mechanically interacting molecules individually. Lastly, we used a more abstract theoretical physics model, which transfers our results to all systems with one-dimensional geometry and local interactions. All three approaches give the same results: local interactions that persist only once they cover the whole system affect smaller systems more strongly; local interactions that are relatively stable even before spreading affect larger systems more strongly.

## Introduction

Changes of behavior in spatially distributed systems are often the consequence of local events that spread to global extent. Examples include nucleation of phases of matter^1^, traffic jams^2^, or sociopolitical phenomena^3^. The global impact of local events depends both on the spreading behavior of such events, as seen for disease outbreak^4^, and the system’s overall geometry, as seen in obstacle-induced symmetry breaking in reconstituted heart tissue^5^. A fundamental question that arises is how the spatial spreading of local events and overall system geometry together determine whether system-wide changes occur.

We wanted to address this question in an example system. Ideally, such a system would undergo frequent and obvious changes in system-wide behavior, which result from well-understood and experimentally adjustable local interactions, and exhibit a range of system geometries. These criteria are fulfilled by the *in vitro* motility assay^6^, where actin filaments of different lengths are propelled by surface-bound groups of myosin molecular motors (Figure 1A). Individual myosin motors interact with equidistant binding sites^7^ along the actin filament (Figure 1B). The resulting unidirectional motion of the actin filament is characterized by periods of arrest and continuous forward sliding (Figure 1C). With increasing filament length, filaments are increasingly biased into the sliding state (Figure 1C). Previous theoretical work explained the observed periods of arrest and sliding by two group kinetic states that emerge by mechanical coupling of myosins via the actin filament^8^ (Figure 1D). Further, it was shown that experimentally induced changes in single myosin mechanochemistry proliferate into altered actin sliding patterns^9,10^. Lastly, statistical distributions of actin sliding velocities for different lengths of filaments can be extracted using automated actin tracking software^9^. Taken together, the *in vitro* motility assay is a good example system to study how locally coupled, experimentally modifiable, and theoretically well-understood myosin kinetics control the bistable switching in the sliding of actin filaments of different length.

**Figure 1:**
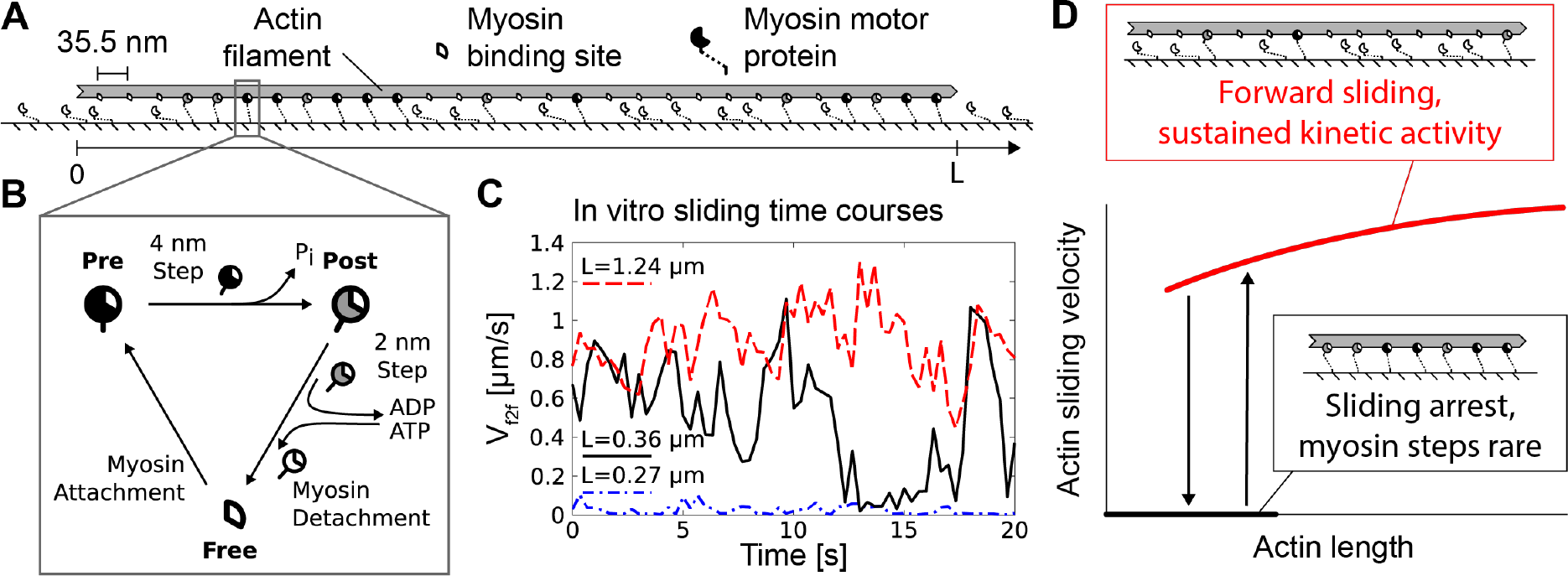
In vitro propulsion of single actin filaments by muscle myosin groups exhibits an actin length dependent stop-and-go pattern. **A)** In our motility assay, purified smooth muscle myosin motors were immobilized on a glass surface to mediate the unidirectional sliding of fluorescently labeled actin filaments. Myosin can bind to actin every 35.5 nm^7^. Myosin binding is oriented with respect to the filament sliding direction. **B)** Individual myosin binding sites go through unidirectional kinetic cycles consisting of three main states^8,10^: unoccupied (Free), pre power stroke (Pre), and post power stroke (Post). The transitions occur by load-independent myosin attachment and load-dependent mechanochemical transitions. **C)** Three example time courses showing instantaneous sliding velocities (“frame-to-frame velocities”, *V*_*f*2*f*_) for actin filaments of different length (*L*). Previous work has shown that actin sliding exhibits *L*-dependent, bistable switching^8,10^; the shortest actin filaments (diffraction-limited, *L* ≲ 0.3 μm) are permanently arrested; slightly longer actin filaments exhibit bistable switching between sliding and arrest; for *L* ≥ 1 μm, actin slides continuously. **D)** Previous work explained how the mechanical coupling of individual myosin motors via the actin filament results in two emergent states in myosin groups, which underlie the actin stop-and-go motion^8,10^. For short actin, myosins bind to actin but fail to effectively undergo mechanical steps due to the mechanical hindrance exerted by other bound myosins. For long actin, myosins bind, execute their mechanical steps, and rapidly unbind again, leading to sustained forward sliding of the actin filament. For actin of intermediate length, alternation between the two emergent states results in a stop-and-go sliding pattern.

Here, we used the *in vitro* motility assay of myosin purified from phasic smooth muscle and performed simulations with single-molecule resolution to investigate how different types of local molecular mechanical perturbations impact the sliding of actin filaments of different length. First, we investigated spontaneous kinetic arrest. Combining experiments and simulations, we found that spontaneous arrest initiates locally, involving a few myosins at close-by positions on the actin filament. Such arrest becomes long-term stable only when it affects further myosins and in this way spreads to reach both ends of the filament. Before the arrest has reached both filament ends, it is readily dissolved. Spontaneous arrest therefore more frequently leads to arrest of shorter actin filaments. To generalize this finding, we implemented an abstract model consisting of a linear spin chain with nearest neighbor coupling. In this abstract model, spontaneous arrest also more frequently spreads to global arrest in shorter chains. Second, we introduced the smooth muscle crosslinker filamin A, which establishes relatively stable actin-surface links throughout the length of actin^11–13^. Again combining experiments and simulations, we found that persistent local kinetic arrest is established once such a link is established, and subsequently spreads to arrest myosin kinetics globally. Because longer actin provides more possibilities for the establishment of such crosslinks, longer actin filaments are more strongly affected. Again, we could generalize this finding using the abstract spin chain model. Taken together, we found that the probability that local perturbations result in global arrest changes differently with system length, depending on whether these perturbations become stable immediately or only once they have spread to span the entire system.

## Results

### Spontaneous kinetic arrest stabilizes after spreading through the entire actin filament

To understand how local interactions between myosins result in the arrest of the entire actin filament, we developed a spatially resolved theoretical model of myosin group kinetics. Several studies have shown coordination in the kinetics of myosin groups that are internally coupled and externally loaded ^14–18^. It has also been proposed that the unloaded actin sliding in *in vitro* motility assays is affected by coupling via the actin filament^8,9,19,20^. Specifically, myosins exert loads on each other, thereby dynamically altering the stepping rates of individual myosins. Note that a widely used model, where myosins independently interact with actin, and actin slides at maximal velocity whenever at least one myosin is attached^21^, is not compatible with such coordinated myosin group kinetics. We therefore used a more recently proposed model for coupled myosin kinetics^8^, which we extended to account for the spatial distribution of protein attachment points along the actin filament as well as the decay of coupling strength with distance along the actin filament (SI Figure 1). The extended model accurately predicted the effect of a reduction of myosin concentration, suggesting that it is a fair representation of our experiments (SI Figure 2A-F). We therefore studied how, in numerical simulations based on this model, interactions of myosin would lead to actin sliding arrest. Simulations of very short actin filaments exhibited extended periods of global kinetic arrest of myosins (Figure 2A). Simulations of intermediate length actin filaments exhibited alternating periods of kinetic arrest and kinetic activity of myosins (Figure 2B). Simulations of long actin filaments did not exhibit periods of global kinetic arrest (Figure 2C). However, local patches of prolonged binding of myosins to the actin filament did occur. When we measured the spatial extent of such arrest patches, we found that longer patches are generally less frequent than shorter patches (Figure 2D). An exception are patches that span the entire length of the actin filament, which occur with an increased frequency. We therefore hypothesized that the relatively short lifetime of arrest patches limits their longitudinal growth - except when the patches grow to the extent of covering the whole actin filament, and become long-term stable. Indeed, when we analyzed the lifetimes of patches that reach none, one, or both ends of the actin filament, we found that prolonged kinetic arrest occurs only when patches reach both ends of the actin filament (Figure 2E). This finding implies that long-lived kinetic arrest is less likely for longer actin filaments, thus explaining the experimental observation that sliding arrest is less likely for longer actin filaments^8–10^ (SI Figure 2A).

**Figure 2:**
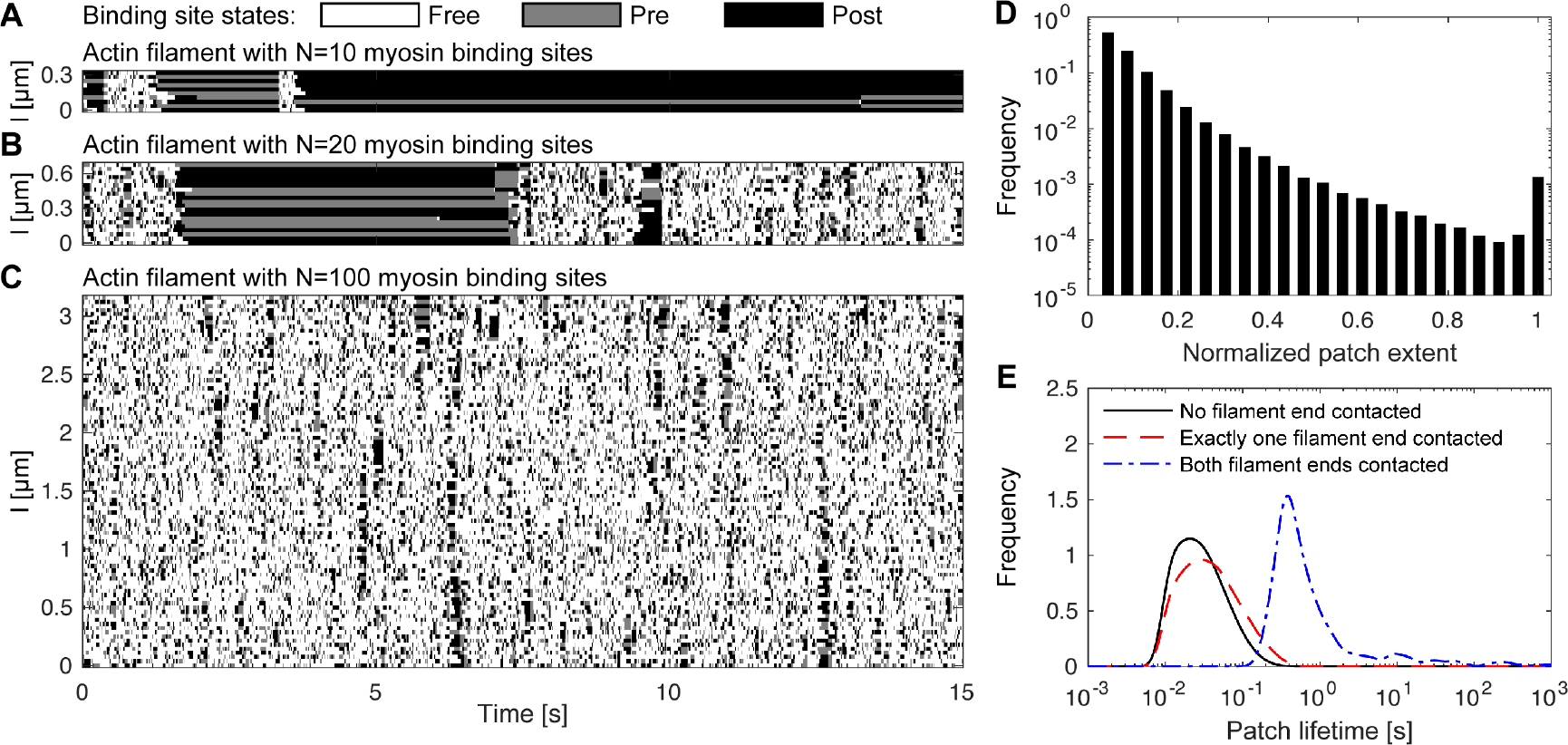
Spontaneous kinetic arrest becomes long-term stable when it spreads to cover the entire actin filament. **A-C)** Example evaluations of our detailed model with single myosin binding site resolution are shown for actin filaments of different length, *L* = *N* × 35.5 nm (*N* refers to the number of myosin binding sites). Myosin binding site states are indicated by color, their position along the actin filament (*l*) by the vertical axis. Evaluations for all *N* exhibited two emergent states: (1) fast cycling through all three binding states, and (2) spatiotemporally contiguous domains of kinetic arrest, marked by Pre and Post state myosin binding sites, in agreement with previous work^8,10^. At low *N*, kinetic arrest dominates (A). At intermediate *N*, kinetic arrest and fast cycling alternate (B). At high *N*, kinetic arrest remains transient (C). **D)** Histogram of the maximal spatial extent of individual arrest patches, normalized by total actin length *L*. In general, patches of larger extent are less frequent. This trend is inverted for patches that extend over the entire actin filament, which are again more frequent. Individual arrest patches were detected as temporally and spatially contiguous regions (connected components) of myosin-occupied binding sites. A single model evaluation over 10,000 s of an intermediate length filament (*N* = 23, or *L* = 816.5 nm) was analyzed so as to cover arrest as well as sliding phases. The model evaluation was resampled at 20 ms intervals for analysis, arrested patches that persisted for 100 ms or longer included in the analysis.) **E)** Empirical probability distributions of patch lifetimes of patches that contact no end, one end, or both ends of the actin filament during their lifetime. (Same model evaluation as panel D, empirical probability distributions obtained by Gaussian kernel estimation with bandwidth 0.1, applied after log_10_transformation.)

### Molecular mechanics can be mapped to a generalized spin chain model

To assess if the conclusions that we made from our example system up to this point can be generalized to other systems, we mapped our findings to a spin chain model (Figure 3A). The spin chain model consists of *N* linearly ordered spins with nearest-neighbor connections. The spins can flip between two states (*s*_*n*=1,…,*N*_ = ±1). This model can be considered representative of the general class of finite, one-dimensional systems with local coupling. The local coupling in our example system is included in the abstract model in the form of a nearest-neighbor coupling with strength *β* > 0. The general bias towards activity exerted by myosin motors is represented by a global field *h* > 0 that acts on all spins. The stabilization of kinetic arrest at the filament ends is represented by a negative field with strength *h*_0_ > 0 that is coupled only to the spins at the chain ends. A numerical evaluation of this model exhibited patches of inactivity, which are similar to the arrest patches seen in our detailed mechanochemical model (Figure 3 B). We adjusted the model parameters (*β, h, h*_0_) against motile fraction (fraction of all observed frames spent in the forward sliding state, *f*_*mot*_) curves of the experimental data set with reduced myosin concentrations (Figure 3C, SI Figure 2G and SI Figure 3). The adjusted model correctly predicted key features of measured autocorrelation time curves (SI Figure 2H), indicating that the spin chain model is a fair representation of our experiment. These findings suggest that results obtained for our mechanochemical example system can indeed be generalized to finite, one-dimensional systems with nearest-neighbor coupling.

**Figure 3:**
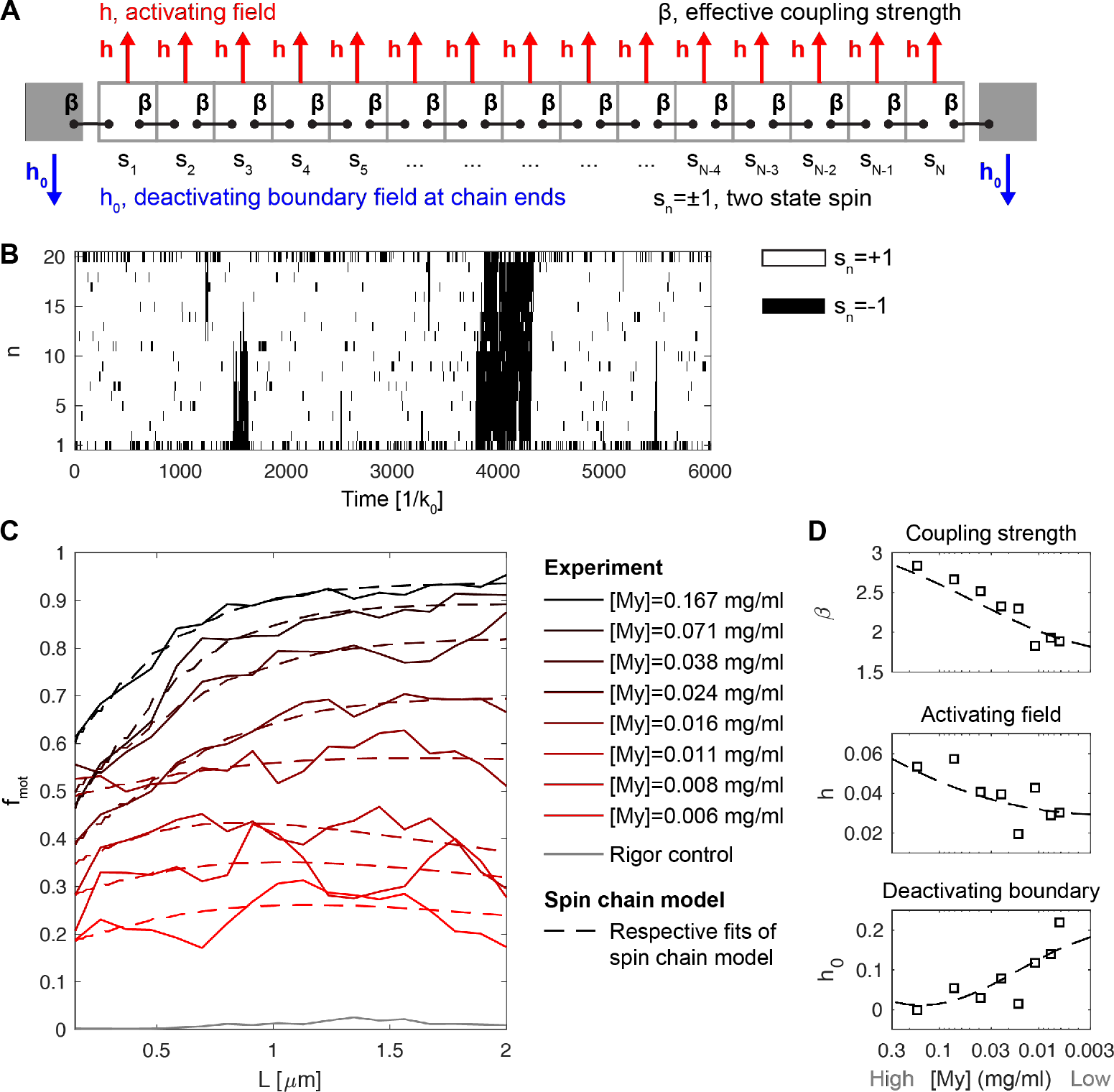
Mapping of molecular mechanics to a generalized spin chain model. **A)**To generalize the behavior seen in our detailed model, we constructed a linear chain of spins with finite length *N*. Spins were either active (*S*_*n*_= +1) or inactive (*S*_*n*_= −1), representing the two emergent kinetic states of the detailed model. Spins were coupled to their nearest neighbors with strength *β* > 0 and were biased towards the active state by a field *h* > 0 affecting all spins in the chain. The spins at the chain ends were coupled to a deactivating field *h*_0_ > 0. **B)** Example evaluation with *N* = 20, active spins are shown in white, inactive spins in black. Model parameters for the highest myosin concentration (0.167 mg/ml) are used. Time is scaled by the characteristic spin flipping rate *K*_0_. **C)** Actin length (*L*) resolved plots of the motile fraction (fraction of total acquired frames that are categorized as actively sliding, *f*_*mot*_) for different myosin concentrations ([My]). To provide permanently arrested actin filaments for reference, a rigor control condition in which myosins attach, but do not detach from actin was included ([My]=0.167 mg/ml, no ATP in the motility buffer). Solid curves are running averages from experimental data (n=8, 9, 9, 8, 8, 9, 9, 5, 3 flow-through chambers per conditions, from top to bottom). Dashed lines are exact *f*_*mot*_ values from the spin chain model fitted to the different conditions. Full experimental data with error bars and corresponding evaluations of both the detailed model and the spin chain model are shown in SI Figure 2. For parameter fitting procedure, see SI Figure 3. **D)**Fitted parameter values (squares) for decreasing [My], with a polynomial fit (dashed line) to guide the eye.

Fitting the generalized spin chain model as described above captures the effect of experimental perturbations in terms of the spin chain model parameters *β*, *h*, and *h*_0_. As an example, let us address the effect of reducing myosin concentrations in our experiments (Figure 3C). The reduced myosin concentration led to a reduction of the bias for kinetic activity (*h* decrease), as was expected (Figure 3D). In addition, the coupling of neighboring domains was compromised (*β* decrease) and the deactivating influence of the chain ends was increased (*h*_0_ increase) (Figure 3D). This mapping to a spin chain model provides an additional theoretical approach to understand how changes in local parameters - such as the effective rate of myosin recruitment - impact system-wide kinetics. While the mechanistically detailed model can reproduce the experimental data starting from the single-molecule level, the spin chain model allows capturing the emergent myosin group kinetics by a set of three effective parameters.

### Kinetic arrest by crosslinkers more strongly affects long actin filaments

To interfere with actin sliding by an external perturbation, we added the crosslinker filamin A to our assay. Filamin mechanically links actin to the motility surface, and forms relatively long-lived bonds (≈ 11 s) unless it is detached by mechanical force mounted by the myosin motors that propel the actin filament (Figure 4A)^13^. We imagine, in principle, two opposing scenarios by which the length of actin could factor into this interaction. (i) As observed above, the kinetically active state that emerges from mechanical coupling of myosins is more robustly preserved for longer actin filaments. This implies that, for longer actin filaments, the actin-bound myosins are more likely to mount the forces required to resolve filamin crosslinks. In this scenario, the sliding of longer actin should be less affected by filamin. (ii) Since mechanical interactions are likely limited within a finite coupling range along the actin filament (SI Figure 1), filamin crosslinks should then also be broken by forces mounted by myosins bound to actin within coupling range. In consequence, the mechanical interaction between crosslinks and myosins can be seen as playing out within a local vicinity. Longer actin filaments would provide more space for such locally confined interactions, each of which has a chance to result in actin sliding arrest. In this scenario, the sliding of longer actin should be affected more strongly by filamin. In our experiments, filamin caused actin to be in the arrested state more frequently, while filamin did not reduce the velocity while actin is actually sliding (see SI Figure 4, SI Figure 5). We therefore focused our further analysis on changes in *f*_*mot*_. For low filamin concentrations ([Fil] ≲ 2.5 nM), *f*_*mot*_ was reduced for all actin lengths, but still exhibited an increase with actin length in agreement with scenario (i) (Figure 4B). For higher [Fil], *f*_*mot*_ was also reduced for all actin lengths, but decreased with actin length in agreement with scenario (ii). We confirmed this second effect in a repeat experiment (SI Figure 6). Also, we found an increased autocorrelation time for intermediate [Fil] (Figure 4C). Taken together with the observation that the arrested and sliding state of actin remain distinct (see SI Figure 4), the increased autocorrelation time suggests that the rate of switching between sliding and arrest was markedly reduced. Our detailed model and our spin chain model reproduced this effect when we introduced crosslinkers that could bind anywhere along the actin filament (Figure 4D-G). Thus, at intermediate to high crosslinker concentrations, perturbations that result from crosslinker binding suppress activity more effectively for longer actin filaments.

**Figure 4:**
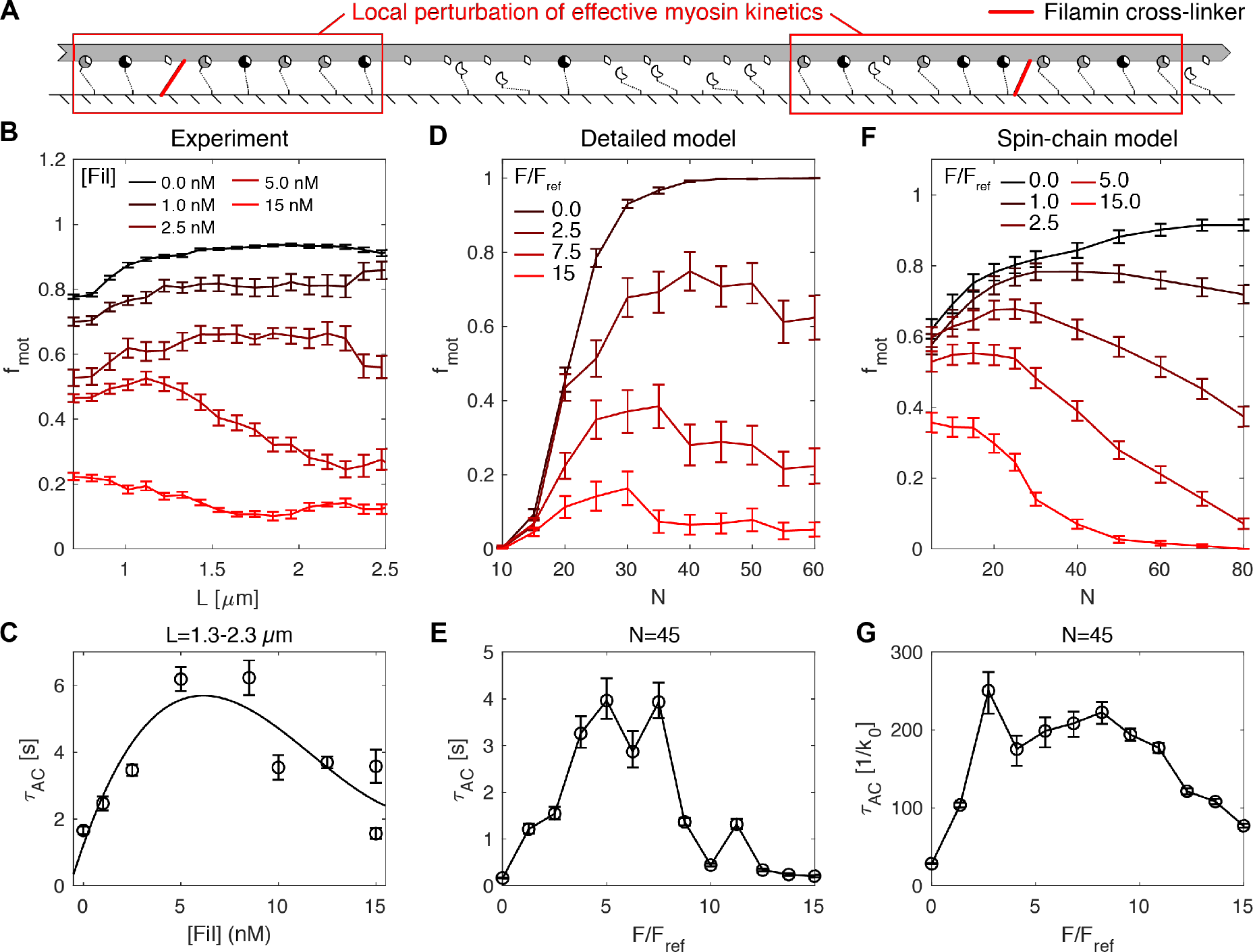
The actin-surface crosslinker filamin more strongly arrests longer actin filaments. **A)** To introduce external perturbations, we added the actin crosslinker filamin to our motility assay. Filamin attaches to actin and the motility surface coating, and is detached by mechanical stress exerted by close-by myosins^13^. **B)** Increasing filamin concentrations ([Fil]) progressively shut down actin sliding. For intermediate [Fil] (5.0 nM), filamin sliding was more effectively stopped at greater actin length (*L*). We confirmed this observation by a second set of experiments (SI Figure 6). Graph shows a running mean±SEM, n=6, 6, 7, 8, 6 flow-through chambers per condition. Note that the analysis of *f*_*mot*_ across the displayed *L* range requires an analysis procedure that obscures the arrest at small *L*, as seen previously^8^. **C)** The sliding velocity autocorrelation time (*τ*_*AC*_) for long actin filaments shows a transient increase for intermediate [Fil], indicating a slower switching between sliding and arrest in this experimental regime. Circles are mean±SEM, n=6, 6, 7, 8, 4, 4, 4, 4, 6 flow-through chambers per condition (data from the second set of experiments were included), left-to-right, top-to-bottom. A cubic polynomial was fitted to guide the eye (solid line). **D-E)** The detailed model reproduced the experimentally observed dependence of *f*_*mot*_ and *τ*_*AC*_ on [Fil] and *L* (*F* is the average number of filamin proteins in binding range per myosin binding site, and is rescaled with *F*_*ref*_, the maximum *F* value used in the model evaluations, which resulted in a good fit to the [Fil]=15 nM condition in our experiments, here *F*_*ref*_ = 0.06/15). All values are mean±SEM, n=32 model evaluations per data point. **F-G**) The extended spin-chain model equally reproduced the experimentally observed dependence of *f*_*mot*_ and *τ*_*AC*_ on [Fil] and *L* (*F*_*ref*_ = 0.3/15). Values are mean±SEM, n=200 evaluations per data point.

**Figure 5:**
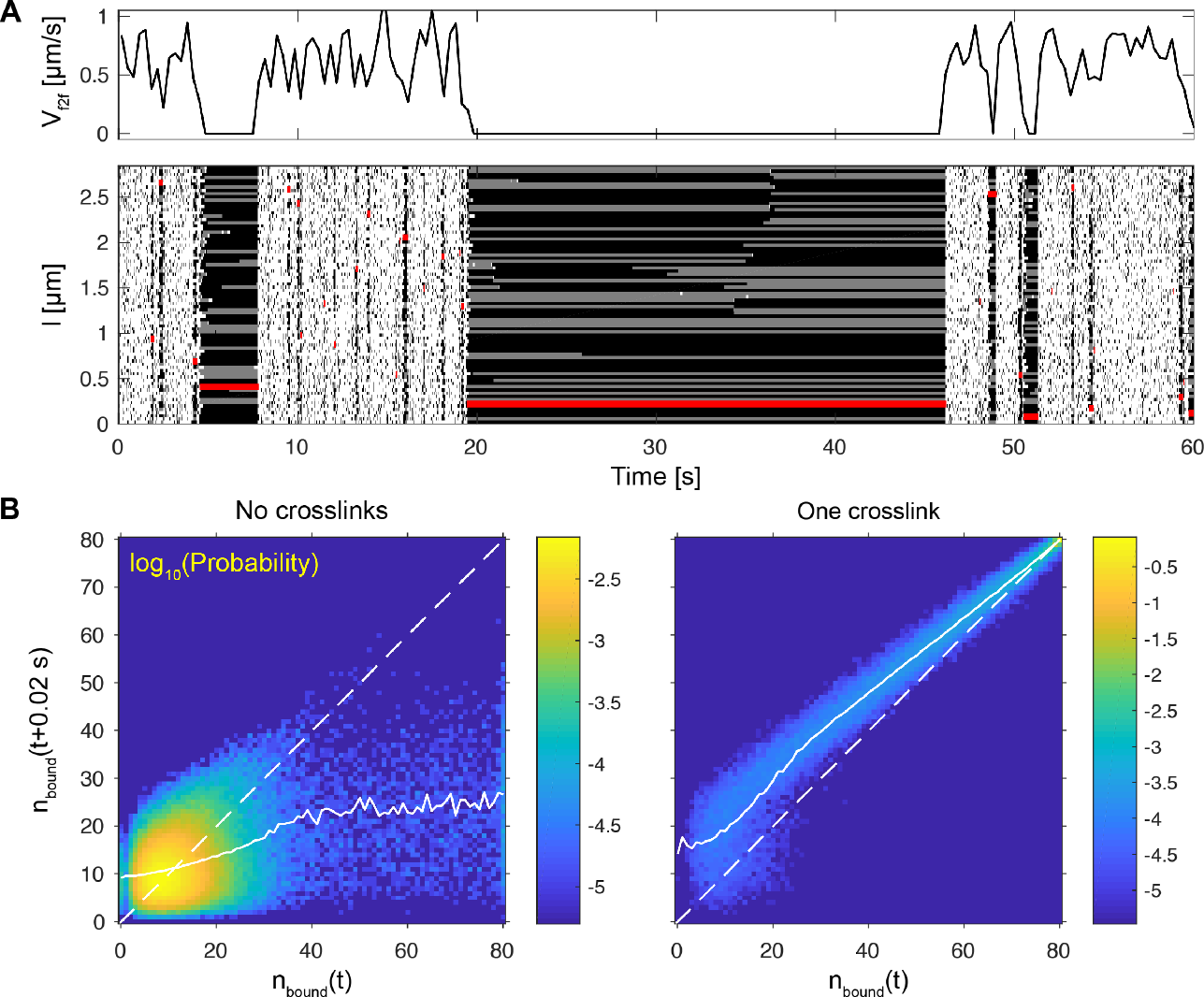
Crosslinker binding allows patches of arrested myosins to spread into global arrest events. **A)** A representative simulation of actin-myosin interaction in the presence of one filamin protein (*N* = 80 myosin binding sites, corresponding to an actin length of *L* = 2.8045 μm, presence of only one filamin protein was assumed to prevent very long-lived actin sliding arrest events). The instantaneous sliding velocity (*V*_*f*2*f*_) time course shows transitions between uninhibited sliding and arrest. The kymograph (see Figure 2 A) shows that myosin arrest and filamin binding (displayed in red) frequently occur at the same time. **B)** Empirical return maps constructed from the number of bound myosins (*n*_*bound*_) at a given time point *t* and with a delay of 0.02 *s* (pooled from 500 simulations of 200 *s*). The average time evolution can be assessed by a comparison of the mean *n*_*bound*_(*t* + 0.02 *s*) for a given *n*_*bound*_(*t*) (solid white line): where the solid white line is above the *x* = *y* diagonal (dashed white line),*n*_*bound*_ increases with time; where it is below, *n*_*bound*_ decreases with time. Points were added to the return maps selectively dependent on no filamin being bound (No crosslinks) or one filamin being bound (One crosslink) at *t* + 0.01 *s*, color map represents the *log*_10_ of the empirical probability.

### Crosslinkers allow patches of arrested myosins to spread into global arrest events

To explain how filamin might exert the observed effects on actin sliding, we investigated simulation results from both our models more closely. In the detailed model, binding of filamin appeared to be correlated with periods of sliding arrest (Figure 5A). To investigate this apparent coordination, we extracted conditional return maps that were gated based on the binding of one filamin protein (Figure 5B). These return maps revealed that, during periods when filamin is bound, the number of bound myosins increases to full occupation of the actin filament. With no filamin bound, the number of bound myosins decreases to a level typical of processive forward sliding of the actin filament. Hence, in the detailed model, the binding of a single filamin to actin is sufficient to stabilize local patches of arrested myosin, which then spread and ultimately cover the entire actin filament. In the spin chain model with added crosslinkers, the binding of crosslinkers similarly allows the spreading of local patches of deactivated spins to cover the entire spin chain, in a manner comparable to the detailed model (SI Figure 7). Thus, both models suggest that the local perturbation by crosslinkers is relatively persistent, and can sustain the spreading of patches of arrest into global arrest. Assuming a constant per-site likelihood of crosslinker binding, larger domains provide more opportunity for such perturbations to effect global kinetic arrest. Our experimental results demonstrate such a case, and our models provide a detailed explanation of the observed increase in actin sliding arrest with actin length.

## Discussion

In this study, we found that local molecular mechanical perturbations impact the sliding of actin filaments differently depending on how these perturbations spread along the actin filament. Perturbations that stabilize only once they have spread throughout the entire filament are less likely to interfere with the sliding of longer filaments. Perturbations that are relatively stable upon their occurrence and subsequently spread into global arrest are more likely to interfere with the sliding of longer filaments. This principle also applies to one-dimensional, finite, and locally coupled systems in general, as demonstrated by our spin chain model.

To explain the sliding patterns of actin filaments that we observed in our experiments, we simulated the interaction of myosin and filamin with the actin filament at single-molecule resolution. Expanding on a previously developed model with single-molecule resolution^8^, we here additionally considered the spatial distribution of protein attachment points along the actin filament. This extended model allowed us to explicitly account for the decay of the strength of mechanical coupling with greater distance along the actin filament. In the previous model, which had no such spatial component, a saturation of the number of maximally coupled myosins had to be assumed *ad hoc* for long actin filaments^8^. Our extended model inherently accounted for the saturation of actin sliding velocity when we included an exponential decay with a characteristic length of ≈ 0.3 μm, and also when we included a linear decay with full loss of coupling at a length of 0.8 μm. We interpret this as indication that the precise shape of the decay is not relevant to explain the observed saturation of actin sliding velocity, and that approximately half of the coupling strength is lost at a range of 300–400 nm. To compare this value to an independent estimate, we calculate the length *l*_*equ*_ at which the longitudinal stiffness of actin 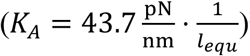^22^ is equal to the compound stiffness of a group of myosins that fully occupy a length of *l*_*equ*_ (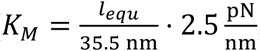, assuming one taut myosin every 35.5 nm)^7,15^. We interpret the resulting *l*_*equ*_ ≈ 130 nm as a lower bound that supports the characteristic length estimate obtained by fitting our model. Taken together, we have now obtained a physically credible, single-molecule level model of *in vitro* actin propulsion, which explicitly treats local mechanical interactions between proteins bound at different positions on the actin filament.

Upon evaluation of our extended model, we observed coordinated behavior of myosin and other proteins, which was spatially distributed along the actin filament. Specifically, we found that the spreading and shrinking of patches of kinetically arrested myosins plays crucial roles in actin sliding and arrest. We are not aware of any existing experimental technique that could record such behavior from actual *in vitro* motility assays. Coming closest, the coordinated conformational changes of actin-regulatory proteins have been resolved in time and along the length of actin filaments by fluorescence microscopy^23,24^. Notably, these experiments revealed patches of activation that initiate locally and then spread along the actin filament, underlining the importance of one-dimensional spreading phenomena in the context of actin-based systems. Also, first experimental successes in tracking several motor proteins within the context of mechanically coupled groups have been reported^23^. Both experimental advances indicate that the spatiotemporal coordination of molecular kinetics along actin filaments might be experimentally accessible in the future. Our models provide highly specific predictions for observations from such experiments, pointing towards the dynamics of patches of arrested myosins as a key factor in *in vitro* actin propulsion.

We proposed an abstracted model consisting of a linear chain of spins, which reproduced key phenomena seen in the detailed model. The close correspondence between the detailed and the abstract model strongly suggests that the conclusions drawn from our experiments and the detailed model can be transferred to any other system that can be abstracted to the same type of spin chain model. This correspondence is therefore a crucial step in transitioning from observations in our particular example system to the generic class of finite, one-dimensional systems. Shortly before submitting this article, we learned that the coil-to-helix transition in model peptides was mapped to a near-identical spin chain model^25^. Specifically, the helix and the coil state correspond to the active and arrested state in our model, respectively. Experimentally, the lifetime of global helix conformations increases with peptide length^25^. Theoretically, this length-dependence was attributed to unfolding that initiates at the peptide ends and is less likely to spread to the center of longer peptides^25,26^. Also, modifications at the peptide ends can influence global outcomes, as found in our work^25^. These experimental observations and theoretical explanations closely correspond to our findings. This correspondence strongly supports our suspicion that our detailed model of actin propulsion maps to the generic class of linear spin chains. Such generic system classes allow to neglect most details of underlying mechanisms. Instead, they capture emergent behaviors whose governing laws apply independently of underlying microscopic processes.

## Methods Summary

### Protein purification and preparation

Phasic smooth muscle myosin and actin were purified from tissue donations (Marvid Poultry, Montréal, QC, Canada), and prepared for motility assays as previously described^27–29^. Purified smooth muscle filamin A was generously provided by Apolinary Sobieszek. Details are described in the SI.

### Motility assay, video recording, and analysis of actin sliding

Buffers, proteins, and flow-through chambers were prepared including several steps for the removal of catalytically defective myosin as previously described^8,10^. Concentrations of myosin and filamin were controlled by dilution in the initial myosin perfusion buffer and the final motility buffer, respectively. Videos of the sliding of fluorescent actin were recorded with a high numerical aperture objective and a CCD camera, and analyzed with our ivma^3^ software suite (available as open source) as previously described^9^. The resulting frame-to-frame velocity (*V*_*f*2*f*_) time traces with an effective time resolution of 0.33 seconds were used to calculate the mean actin sliding velocity (*v*), the motile fraction (*f*_*mot*_), and the ensemble-corrected autocorrelation time (*τ*_*AC*_, SI Figure 8) for different actin lengths (*L*). Details are described in the SI.

### Detailed model of actin, myosin, and filamin interaction

We simulated the mechanochemistry of individual myosin binding sites on a given actin filament as well as filamins crosslinking actin and the motility surface. To introduce the decay of mechanical coupling between proteins more distally placed along the actin filament, we rooted this mechanochemistry in local mechanical equilibria, evaluated from the location *x* of a given protein on the actin filament,

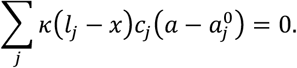

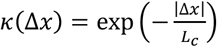 represents a decay of mechanical coupling strength along the actin filament, with a characteristic length *L*_*c*_ = 0.3 μm. *l*_*j*_ indicates the attachment point of a given protein *j* on the actin filament, *c*_*j*_ the stiffness, and 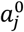 the linear sliding position of the actin filament at which the protein ã is unstrained. The assumption of quasi-instantaneous mechanical equilibration is supported by previously measured longitudinal deformation response times of actin, which are well below 0.2 ms^22^. Further, consideration of only the linear sliding without viscous drag from the surrounding solvent is well-justified in the motility assay as we prepared it^21^. The local mechanical equilibrium is attained by adjusting the sliding position *a* of the actin filament. All reactions were assigned reaction rates without mechanical load, and reactions that included mechanical steps were additionally scaled by a prefactor 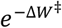 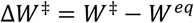 is the mechanical work required to reach the mechanically strained transition state (*W*^‡^) starting from the local mechanical equilibrium state (*W*^*eq*^). The mechanical work levels *W*^‡^ and *W*^*eq*^ can be calculated by applying the local mechanical equilibrium condition with and without the additional strain for the protein of interest to reach its transition state, respectively. The resulting rates were included in an extended Gillespie algorithm, to simulate actin sliding, which was sampled at time intervals Δ*t* = 0.33 s as described previously^8,10^. Details are described in the SI.

### Spin chain model

We implemented a linear chain of spins, *s* =(*s*_1_, *s*_2_,…, *s*_*N*_), where *s*_*n*_ = ±1. The spin chain potential energy was

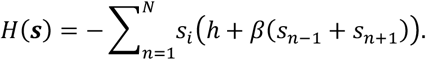

Here *h*>0 is an activating bias affecting all spins and *β* > 0 the connection strength between nearest neighbors. The ends of the spin chain were padded by *s*_0_ = *s*_*N*+1;_ = −*h*_0_, so that *h*_0_ > 0 described a deactivating bias at the ends of the chain. From *H*(***S***), the probability that all *S*_*n*_ = 1 could be exactly calculated to determine *f*_*mot*_. Dynamics of the spin chain were analyzed using a Gillespie algorithm implementation. The effect of crosslinkers was added by a second variable *c*_*n*_ ∈ {0,1}, which monitors the binding of crosslinkers. Spins and crosslinkers influenced each other’s kinetic rates, and their kinetics were also evaluated using the Gillespie algorithm. Details are described in the SI.

## Acknowledgments

We acknowledge funding from the Helmholtz Program Biointerfaces in Technology and Medicine (BIFTM), the Max Planck Society, and the Natural Sciences and Engineering Research Council of Canada (NSERC). The Meakins-Christie Laboratories (McGill University Health Centre) are supported in part by a center grant from Le Fonds de la Recherche en Santé du Québec. Z.B. was supported by the Costello Fund and a Master’s Training Award from Le Fonds de la Recherche en Santé du Québec. L.H. was supported in part by an ELBE Postdoctoral Fellowship from the Center for Systems Biology Dresden. We thank Marvid Poultry for tissue donations; Apolinary Sobieszek for purified filamin; Pradeep Kumar Mohanty for discussing the spin chain model; Patrick McCall and Daniel Riveline for manuscript comments; Tyler Harmon, Alf Mansson, Thomas Quail, Sam Walcott, and Christoph Weber for discussions; and the Max Planck Institute for the Physics of Complex Systems for computing resources.

## Supplementary Materials

**SI Figure 1:**
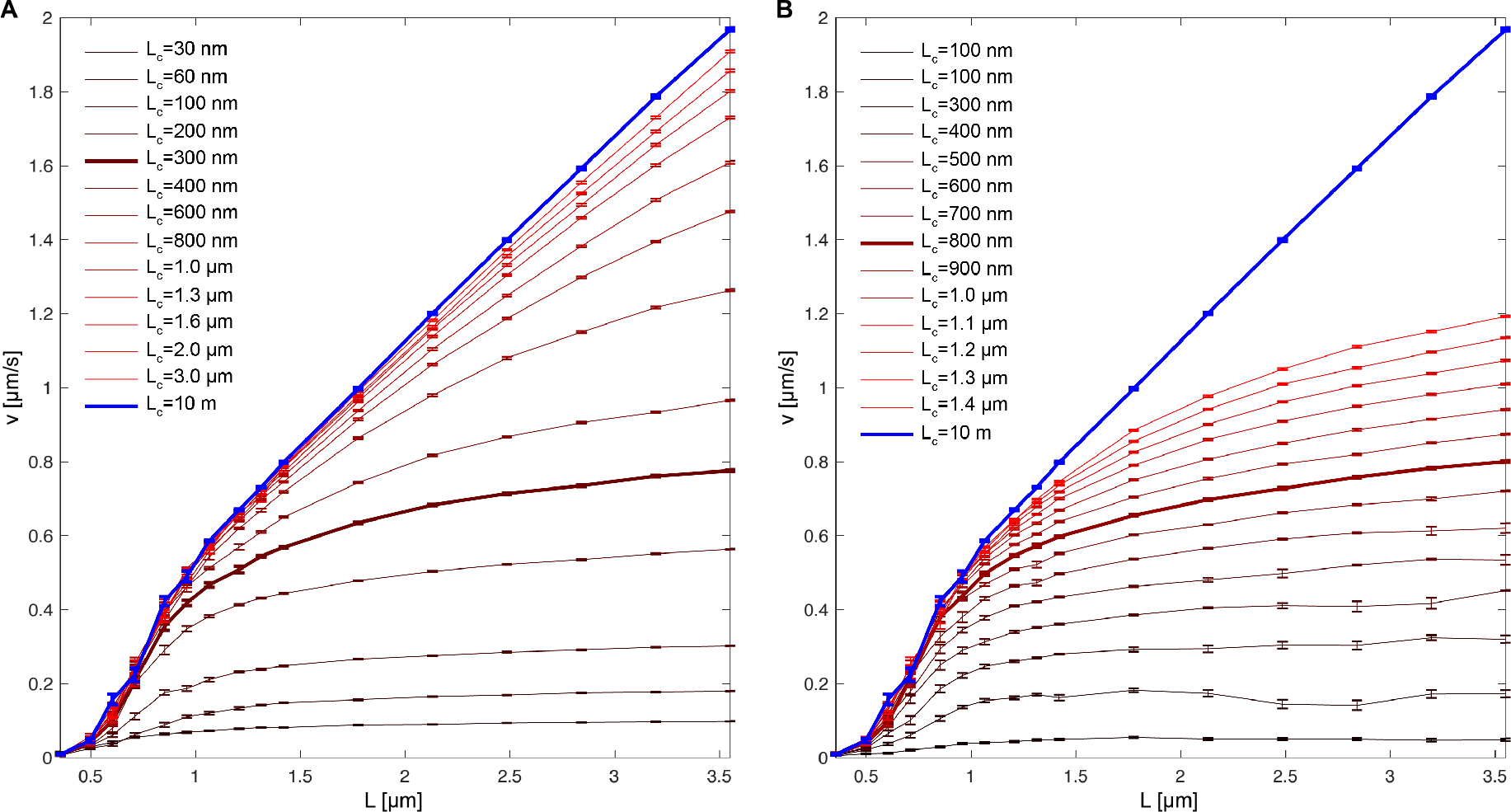
A limited range of mechanical coupling along the actin filament results in more realistic simulation results. For a more realistic coupling between myosins interacting over the length of the actin filament, we introduced a finite coupling length (*L*_*c*_). **A)** We implemented an exponential decay of the coupling strength along the actin filament. We tested different values for the characteristic exponential decay length (*L*_*c*_). A value of *L*_*c*_ = 0.3 μm resulted in a plateau velocity of ≈ 0.8 μm/s, which agrees with the velocity observed in previous experiments^9,10^. We therefore used *L*_*c*_ = 0.3 μm in our simulations. **B)** A linear decay in coupling strength equally limited the plateau velocity, indicating that the specific shape of the decay function is not of primary importance. In the linear implementation, *L*_*c*_ indicates the distance at which the coupling strength reaches zero. In the case of a linear decay, *L*_*c*_ = 800 nm resulted in a plateau close to 0.8 μm/s. Graphs show mean±SEM, 40 simulations per data point.

**SI Figure 2:**
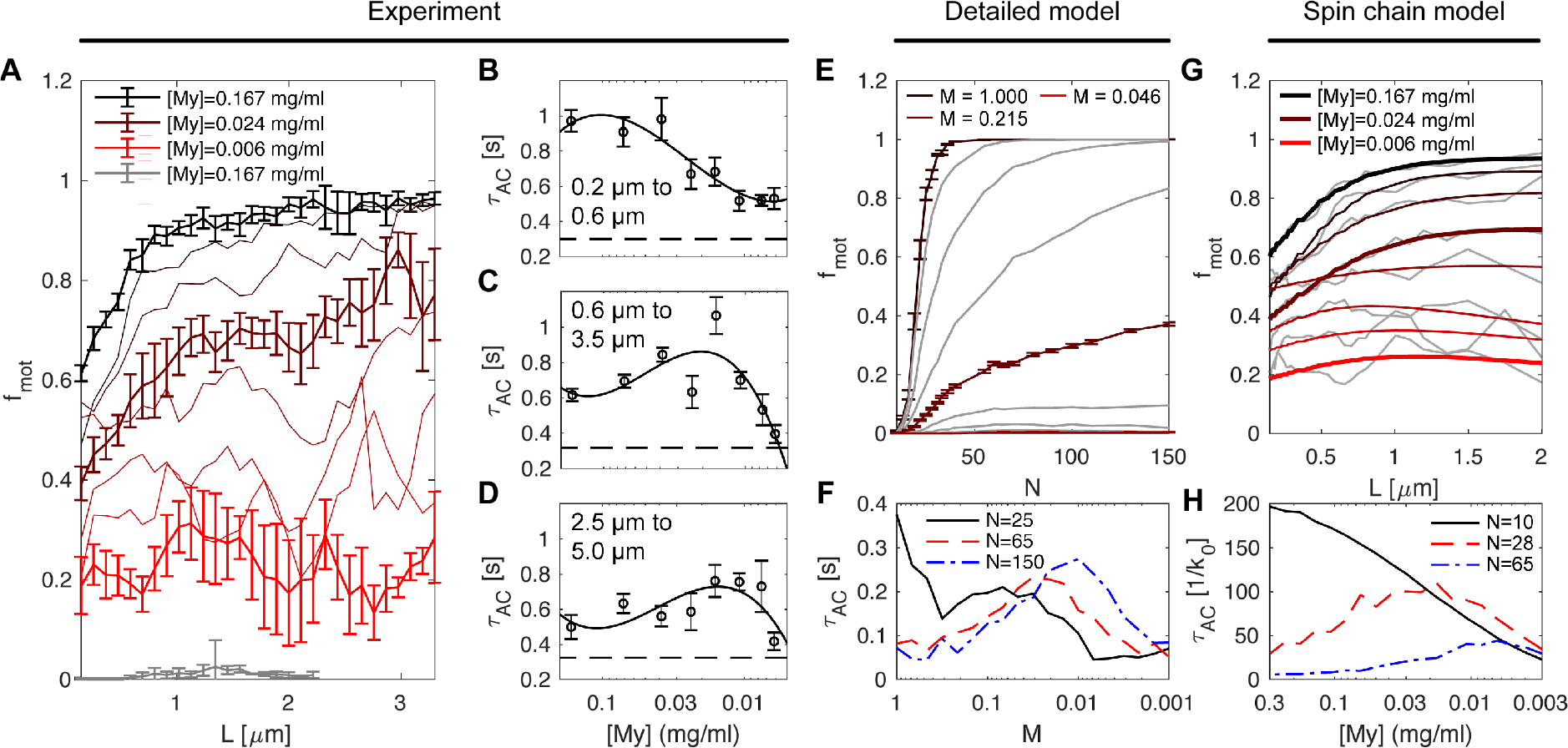
The detailed model and the spin chain model predict the effect of reduced myosin concentrations. To assess whether our theoretical models have predictive power with respect to our experiments, we compared simulation results to experiments with reduced concentrations of myosin ([My]). **A)** Reducing [My] in the experiment lowered the fraction of total tracked frames during which actin filaments are sliding (“motile fraction”, *f*_*mot*_). Rigor myosin (no ATP in buffer) was included as a negative control, which was expected to show *f*_*mot*_ = 0, as was the case. *f*_*mot*_ was resolved by *L* using a sliding window average (values are mean±SEM, n=8, 9, 9, 8, 8, 9, 9, 5, 3 flow-through chambers per conditions, from top to bottom). **B)** The autocorrelation time (*τ*_*AC*_) of actin sliding time courses (*V*_*f*2*f*_) showed a maximum when resolved by [My]. The *τ*_*AC*_ maximum for small *L* was at high [My] (values are mean±SEM). **C)** For intermediate *L*, the *τ*_*AC*_ maximum was shifted to intermediate [My]. **D)** For high *L*, the *τ*_*AC*_ maximum occurred for low [My]. **E, F)** The detailed model reproduces key features of the dependence of *f*_*mot*_ (E) and *τ*_*AC*_ (F) on [My] and *L* (values are mean±SEM, n=48 evaluations per [My]). **G)** We adjusted the model parameters of our spin-chain model, *β*, *h*, and *h*_0_, to fit experimental *f*_*mot*_ (light gray lines, for fitting procedures see SI Figure 3). **H)** Using the same spin-chain model parameters, key features of the dependence of *τ*_*AC*_ on *L* and [My] were correctly predicted.

**SI Figure 3:**
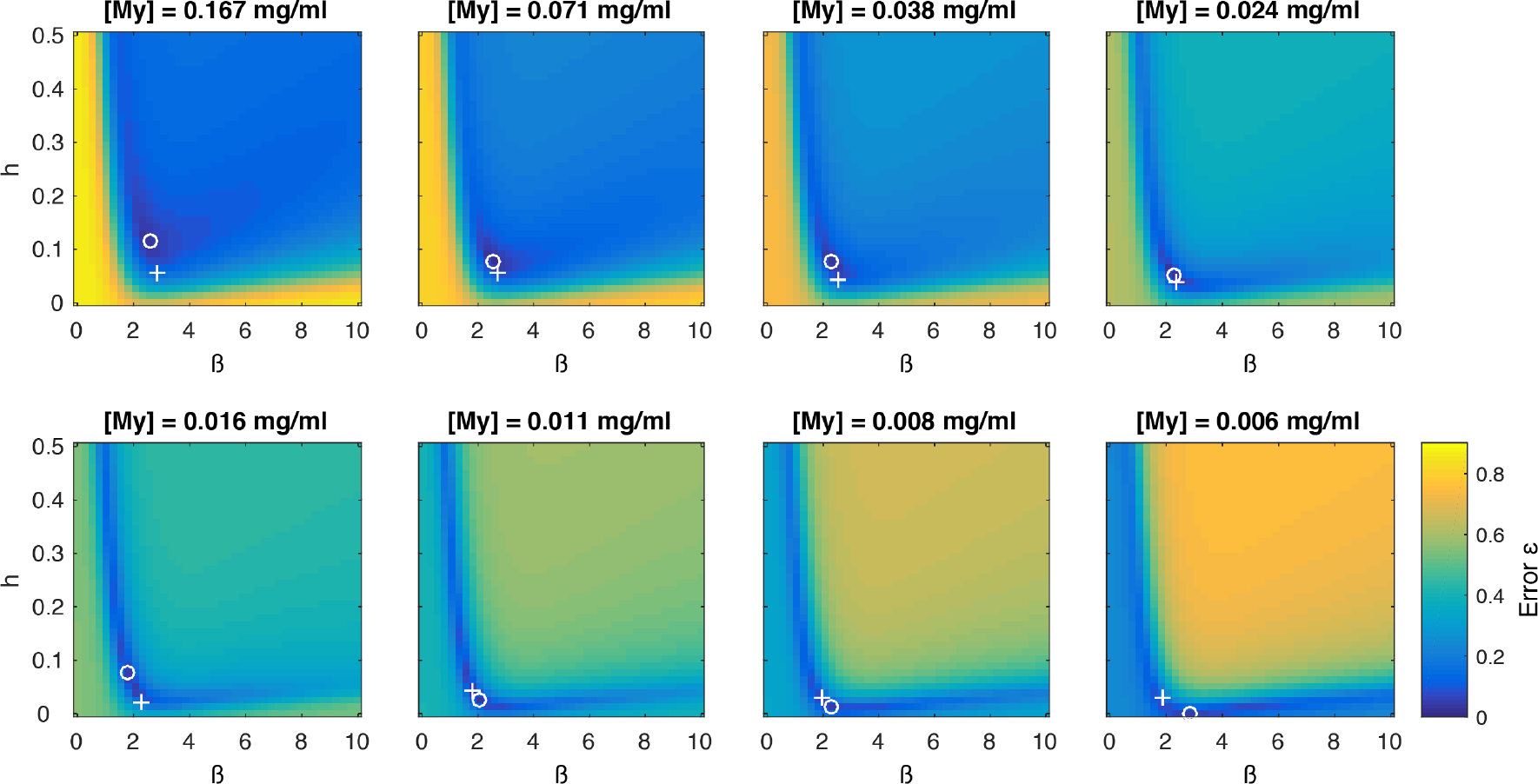
Parameter estimation for spin chain model. The spin chain model was fitted to the motile fraction data for different myosin concentrations ([My]), see Figure 3. The error (*ε*) was quantified as the root of the sum of weighted squared averages. As a first optimization step, *ε* surfaces for the different [My] were calculated (shown as image plots), where *β* and *h* were varied and *h*_0_ = 0.1 was kept constant. The minimum of a given *ε* surface (white circle) was used as the starting point for a second optimization (minimization of *ε*) step, this time without constraining *h*_0_ (white cross). The resulting (*β*, *h*, *h*_0_) combinations were used as parameter sets representing the different [My].

**SI Figure 4:**
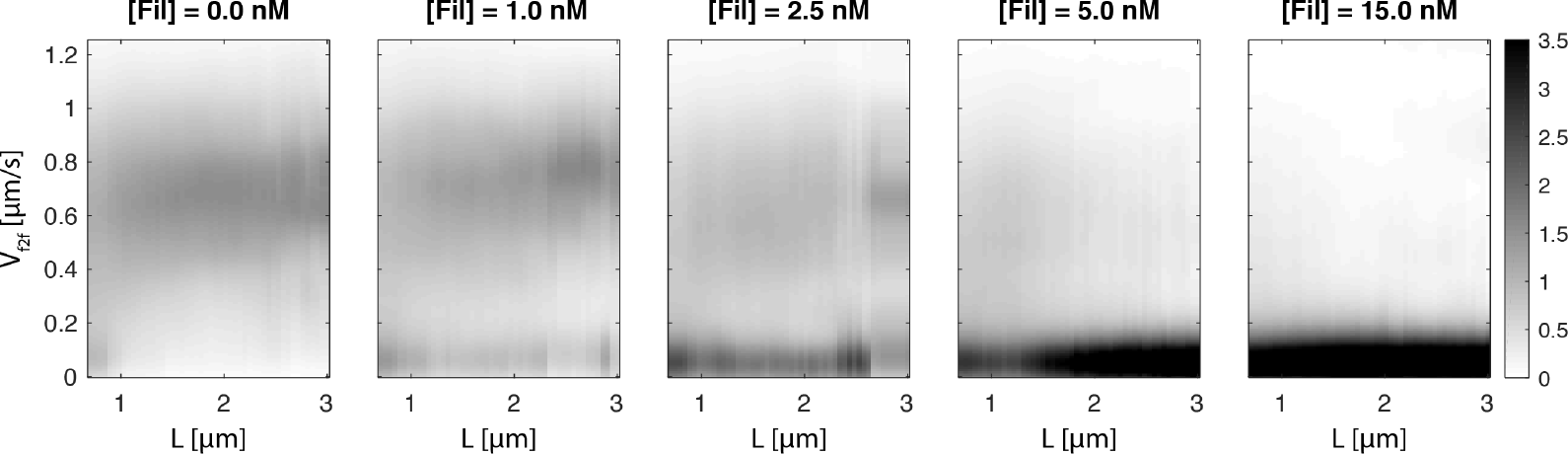
At increasing filamin concentrations, actin is biased into the arrested states, but does not significantly lower the velocity during periods of forward sliding. Panels show experimentally recorded actin sliding velocity distributions for increasing filamin concentrations ([Fil]). With increasing [Fil], the probability to observe the active sliding state becomes less. The velocity of the sliding state is clearly distinguishable for all conditions except [Fil]=15.0 nM, where no more sliding was visible. To create the displayed intensity images, actin filaments were binned by length (*L*), and instantaneous velocities (frame-to-frame centroid displacements, *V*_*f*2*f*_) were used to create empirical probability density distributions at a given length. The intensity of black coloring is proportional to the probability density.

**SI Figure 5:**
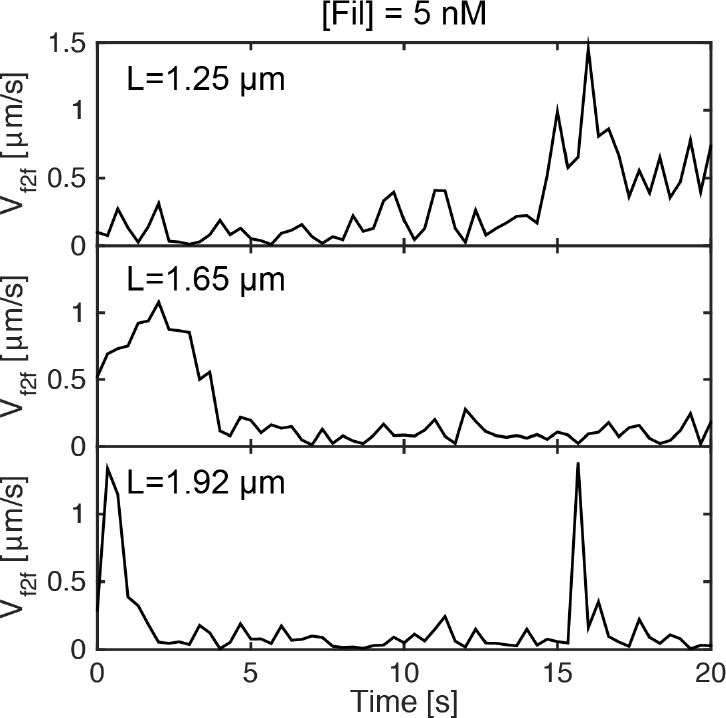
Example traces of actin sliding in the presence of filamin at an intermediate concentration. Example traces were chosen from experiments with intermediate filamin concentration (5 nM) for actin filaments longer than 1 μm to illustrate that filamin halts actin sliding for extended periods of time.

**SI Figure 6:**
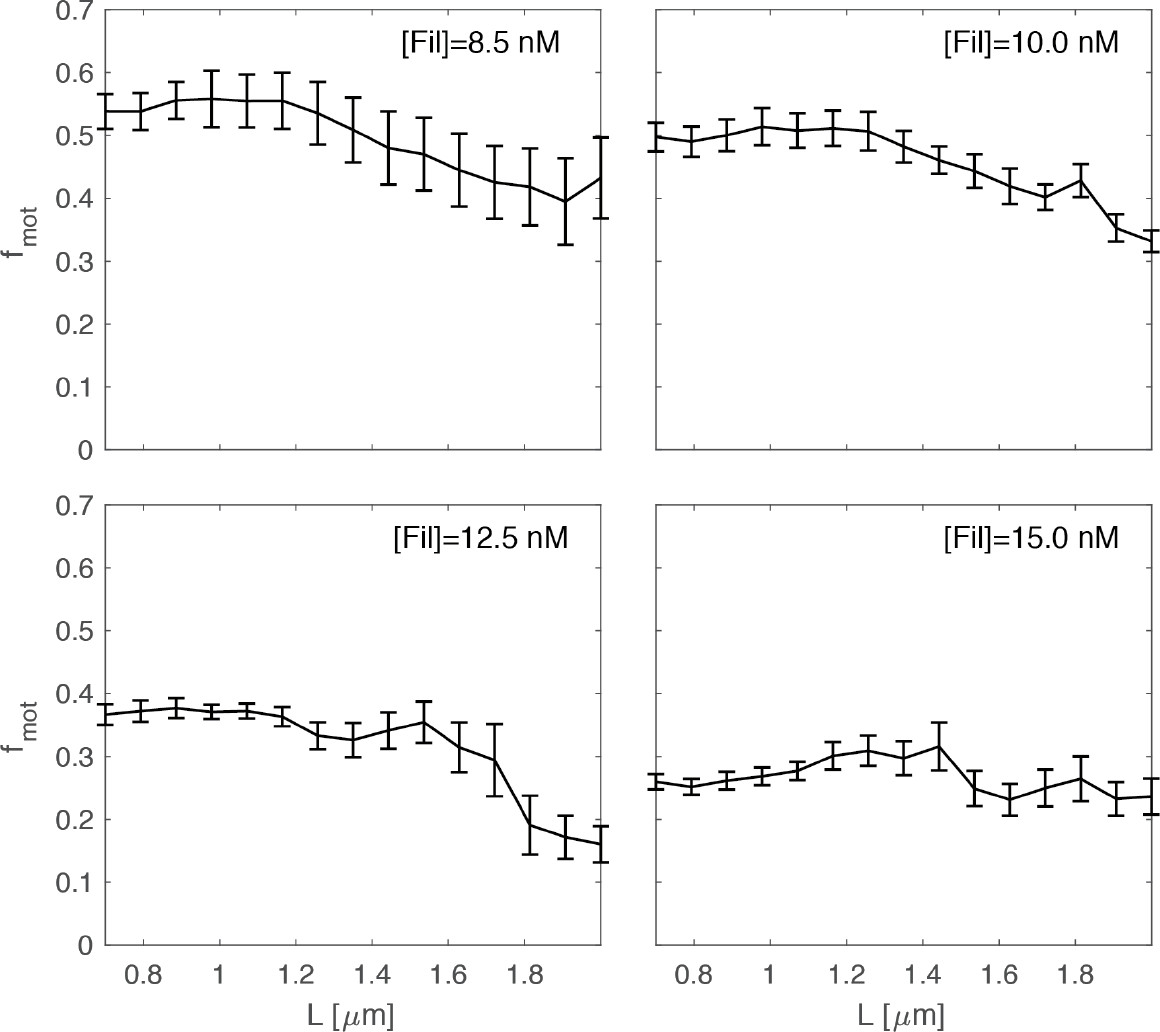
Additional experiments showing that intermediate concentrations of filamin arrest actin sliding more frequently for longer actin filaments. We found that, at intermediate filamin concentrations, the inhibition of actin sliding was more pronounced for longer actin filaments (Figure 4B). To confirm this assessment, we carried out a second set of experiments in an intermediate to high filamin concentration ([Fil]) range. As observed in our initial experiments, inhibition of actin sliding is more pronounced for longer actin filaments, at intermediate [Fil]. Curves show the running average of the motile fraction (*f*_*mot*_) for different actin lengths (*L*); mean±SEM, n=4 samples per condition.

**SI Figure 7:**
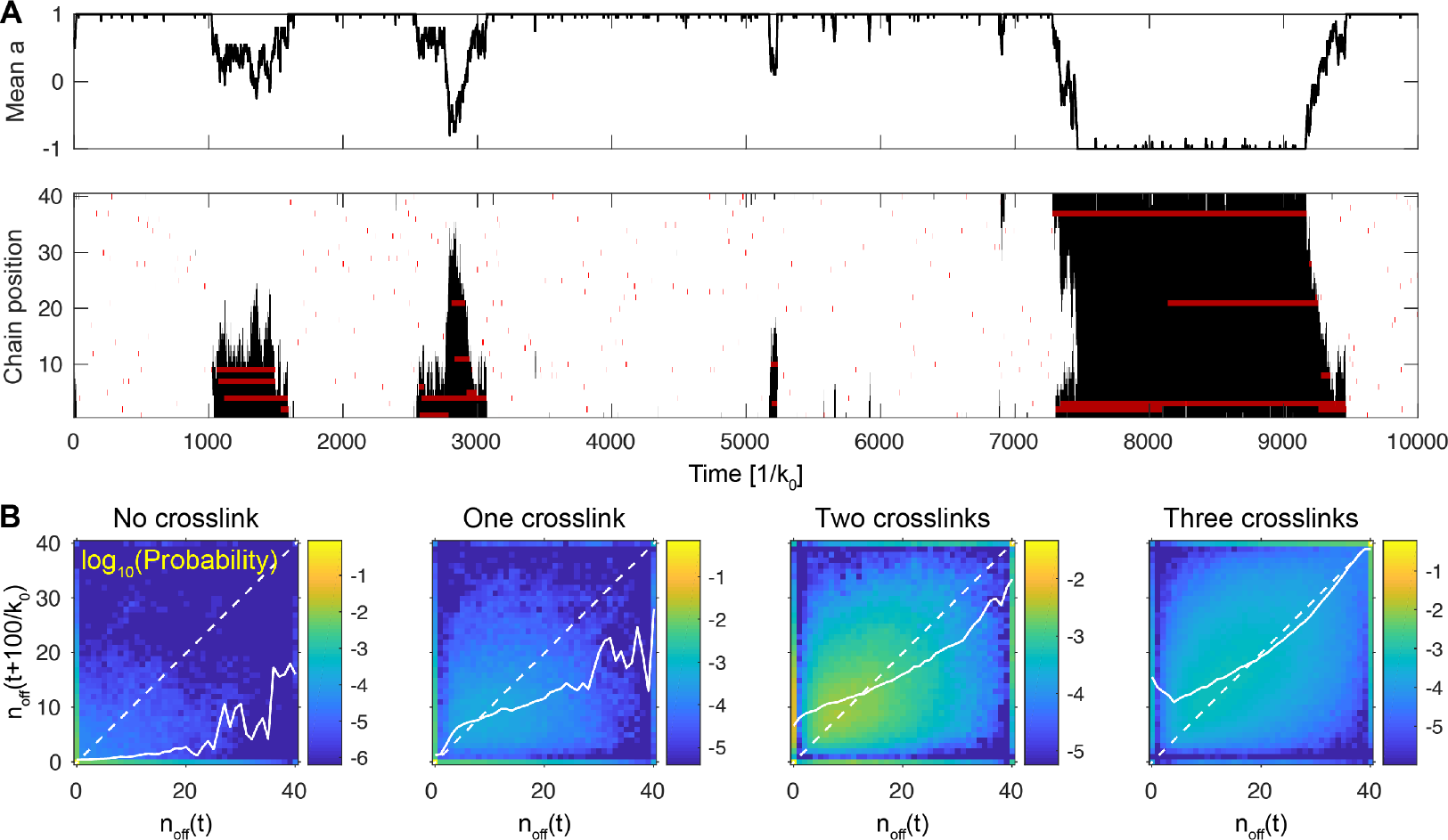
Crosslinker binding stabilizes deactivated domains in the spin chain model. **A)** A representative evaluation of the spin-chain model with crosslinkers. Spins that are on (σ = +1) are indicated in white, spins that are off (σ = −1) in black. Crosslinker binding is indicated by red shading. **B)** Return maps constructed from 200 evaluations of duration 15,000/*k*_0_. The *x* = *y* diagonal is drawn as a dashed white line; the mean *n*_*off*_(*t* + 100/*k*_0_) value for a given *n*_*off*_(*t*) is drawn as a solid white line.

**SI Figure 8:**
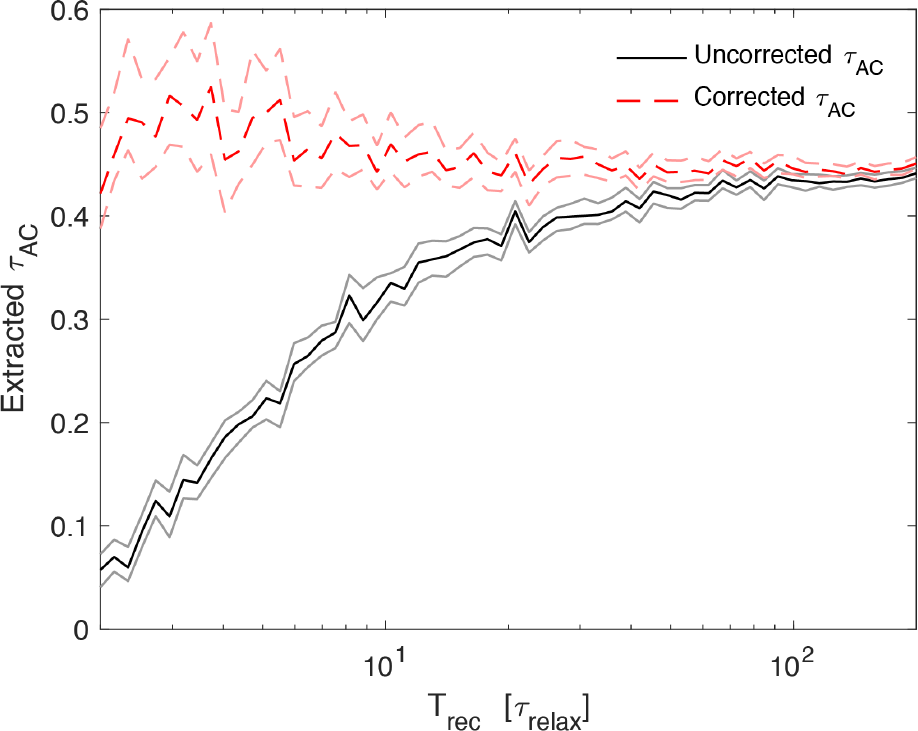
Correction of autocorrelation time calculation for short traces, demonstrated using a bistable toy model system. For dynamic processes that exhibit non-zero autocorrelation, too short recording times (*T*_*rec*_) can yield autocorrelation times (*τ*_*AC*_) that underestimate the autocorrelation time of the actual process^30,31^. We use a toy model simulation with random fluctuations that decay with a relaxation time *τ*_*relax*_ = 0.5 to demonstrate this effect, along with a previously suggested approach for its correction. Applying an uncorrected measure of autocorrelation, *τ*_*AC*_ converged only for *T*_*rec*_ that exceed *T*_*rec*_ ≈ 100 × *τ*_*relax*_ (note that the values of *τ*_*relax*_ and *τ*_*AC*_ do not have to be identical). Applying a corrected measure of *τ*_*AC*_ that considers the ensemble mean over several recorded traces, *τ*_*AC*_ converged already for *T*_*rec*_ ≈ 10*τ*_*relax*_. Data points are mean±SEM, 500 simulation repeats per data point.

### 1 Experimental Methods and Materials

#### Purification of Proteins

Donated tissues from the slaughterhouse (Marvid Poultry, Montréal, QC, Canada) were used for purification. Phasic smooth muscle myosin was purified from chicken gizzards following Sobieszek [1]. Actin was purified from chicken pectoralis acetone powder and stored at 4°C, see Pardee and Spudich [2]. Actin was flourescently labelled by incubation with tetramethylrhodamine isothiocyanate-phalloidin (TRITC P1951; Sigma-Aldrich, Oakville, Ontario, Canada) [3].

#### Myosin thiophosphorylation

Thiophosphorylation of myosin (5 mg/mL) was executed with CaCl_2_ (6.75 mM), calmodulin (3.75 mM, P2277 Sigma-Aldrich), myosin light chain kinase (0.08 mM), MgCl_2_ (10 mM), and ATP γ-S (5 mM). With all the reagents added, myosin was incubated at room temperature for 20 min, kept overnight at 4°C, and then stored in glycerol at −20°C.

#### In vitro motility assay

Myosin was ultra-centrifuged to remove non-functional myosin (42,000 rpm, 4°C, 31 min; 42.2 Ti rotor in Optima L-90K ultracentrifuge; Beckman Coulter, Indianapolis, IN); motility flow-through chambers and buffers were prepared and used as previously described [4]. The oxygen scavenger consisted of 0.25 mg/mL glucose oxidase, 0.045 mg/mL catalase, and 5.75 mg/mL glucose (pH adjusted to 7.4). The motility buffer consisted of actin buffer with additional 0.5% methylcellulose and 2 mM ATP. Myosin stock was diluted to 0.17 mg/mL, or lower myosin concentrations where indicated. Filamin stock (kindly provided by Apolinary Sobieszek) was initially diluted in actin buffer to 4 *μ*M. Filamin concentration was then adjusted by dilution in motility buffer, which was then used in the last perfusion step before video recording. Flow-through chambers were heated to 30°C prior to and during the recording of video data.

#### Video recording

An inverted microscope (IX70; Olympus, Tokyo, Japan) with an oil immersion objective (100x ACH, NA 1.25; Olympus) was used to record actin filament motion. Images were recorded with an image-intensified charge-coupled device camera (30 fps, KP-E500; Hitachi, Tokyo, Japan) connected to a custom-built recording computer (Pinnacle Studio DV/AV V.9 PCI capture card; Norbec Communication, Montreal, Quebec, Canada). Three 30 second long videos were recorded per flow-through chamber.

#### Analysis of actin sliding

We used our in vitro motility assay automated analysis (ivma^3^, available as open source repository on GitHub) to extract instantaneous actin sliding velocity (*V*_*f*2*f*_) traces from our videos [5]. Filaments were individually tracked, sorted by actin length (*L*), and quality control of filament and trace images was carried out using machine learning on manually scored training data. For *L*-resolved plots and statistics, we used sliding window averaging. To prevent artifacts resulting from curved actin sliding traces, only traces with a solidity of 0.5 or greater were included in the analysis. The motile fraction (*f*_*mot*_) was extracted using a *V*_*f*2*f*_ threshold of 0.25 *μ*m/s.

### 2 Ensemble-corrected autocorrelation time

The autocorrelation function (AC) of scalar time series (*x*(*t*)) is a standard approach to determine how strongly fluctuations of *x* at a given time point *t* affect the value of *x* at a later time point *t* + *τ*. The autocorrelation function is commonly defined as

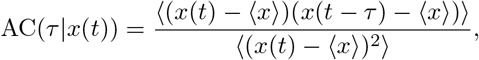

where 〈…〉 indicates a time average. The direct calculation of AC(*τ*) results in too low values unless signal traces are used that significantly exceed the autocorrelation time (*τ*_*AC*_) of a given signal [6, 7]. When several measurements of the same process are available - an ensemble of measurements ({*x*_*n*_(*t*)}, *n* representing different traces from the same process) - a correction can be applied by subtracting not the individual trace mean, but the mean of the ensemble of measurements (*x̄*) [6]. The corrected formulation is

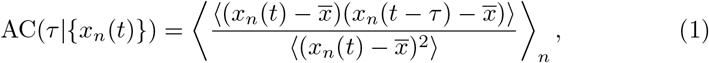

which was applied to all measurements of AC(*τ*).

To determine the average time of the persistence of fluctuations - the autocorrelation time - the lag 1 autocorrelation (*AC*(*τ*=Δ*t*|{*x*(*t*)}), Δ*t* time resolution of time series *x*(*t*)) was used. The lag 1 autocorrelation time was then calculated as

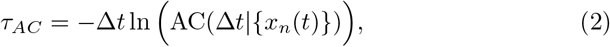

based on the assumption of an exponential decay of the autocorrelation function, 

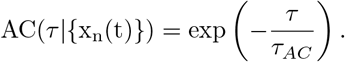

### 3 Detailed Simulations

#### 3.1 Mechanochemistry of myosin and filamin proteins that interact with actin

The interaction of myosin and filamin proteins with a single actin filament of length *L* was simulated. In the case of myosin, this simulation was carried out from the perspective of major myosin binding sites, which are distributed at a fixed distance of *b* = 35.5 nm along the length of a given actin filament, so that the total number of myosin binding sites is *N*_*m*_ = *L/b* [8].

For filamin, the simulation did not include specific attachment points, but the entire actin filament was accessible for binding. To calculate the number of filamin proteins in range of a given actin filament (*N*_*f*_) considering both *L* and differences in filamin concentration, we introduced *F* ∈ [0, 1] as a proxy for filamin concentration. We then calculated

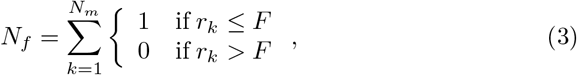

where *r*_*k*_ ∈ [0, 1] are samples from a uniform random distribution. Thus, *F* ∈ [0, 1] represents the probability of a filamin protein being within binding range, with greater *F* representing higher filamin concentrations.

To simulate the linear sliding motion of a single actin filament, the mechanical interactions with myosin motors and filamin crosslinkers were simulated. For both myosin and filamin the principles used to describe the mechanical interaction were identical. For all mechanical calculations, a mechanical equilibrium was assumed to be attained quasi-instantaneously at all times

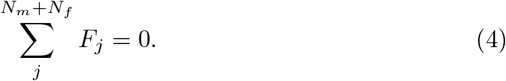

Here *F*_*j*_ is the mechanical force upon the actin filament, mounted by a binding protein *j. j* addresses all *N*_*m*_ myosin binding sites and *N*_*f*_ filamin proteins.

To consider the linear spatial extension of the actin filament, each interacting protein was considered to be attached at a point 0≤*l*_*j*_≤*L* along the actin filament. These attachment points were used to calculate *F_j_* from a reference point 0≤*x*≤*L*, leading to an independent calculation of mechanical equilibria for each individual protein *j*, using *x* = *l*_*j*_. These local mechanical equilibria were described by a modification of Equ. 4,

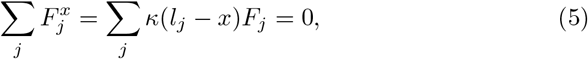

where 0≤*κ*(Δ*x*)≤1 with *κ*(−Δ*x*) = *κ*(+Δ*x*) is a function describing the decay of mechanical coupling with increasing distance along the actin filament (Δ*x*).

Having specified the condition for local mechanical equilibrium (Equ. 5), we will now proceed to satisfy this condition by means of displacing actin in its axial direction. This was formally done by adjusting the sliding position of the entire actin filament (*a*). Any connected protein *j* was assigned a resting position 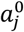, which is the position that actin would slide to if only this protein alone was attached to actin. Note that this resting position can be treated and assigned entirely independently from *l*_*j*_; 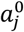 refers to a sliding position of the overall actin filament, *l*_*j*_ refers to a position on the actin filament and uses the actin filament itself as a reference frame. We approximated the force response of all proteins as linear springs,

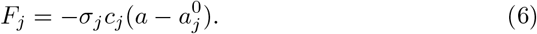

In other words, we assumed that attached proteins are always in the taut chain configuration. Here σ_*j*_ = 1 or σ_*j*_ = 0 indicate that the myosin binding site/filamin protein referred to by the index *j* is attached or not, respectively. *c*_*j*_ is the effective spring constant. For a given protein *j*, the local mechanical equilibrium condition (Equ. 5) always has one unique solution,

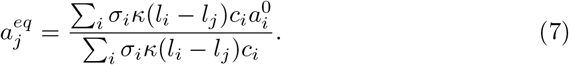

The exception where all σ*_i_* = 0 is circumvented because 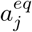 only needs to be calculated when σ*_j_* = 1. Based on Equ. 7, the mechanical work 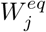 can then be calculated,

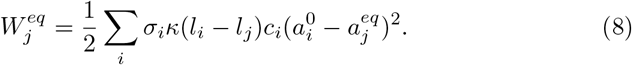

We assigned the decay function

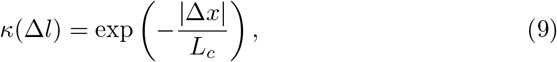

where *L*_*c*_ is the characteristic coupling length along the actin filament. An exponentially decaying coupling strength resulted in good agreement with our experimental data (SI Fig. 1A). The same was true for a linear decay function

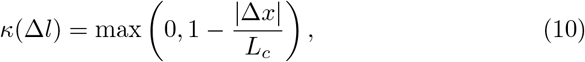

see SI Fig. 1B. Because it seems more reasonable that mechanical coupling decays exponentially in a linear chain of coupled elements, we chose the exponential decay function for further use.

We have now described how to calculate local equilibria, and will proceed to calculate the mechanical work associated with a conformational change 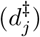 that alters the resting position of a given protein *j*. We captured such a conformational change by an altered resting position 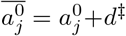. For this conformational change, a new local equilibrium

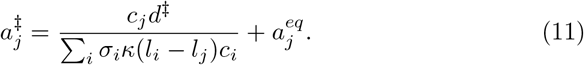

would be attained. The mechanical work associated with this local equilibrium would be

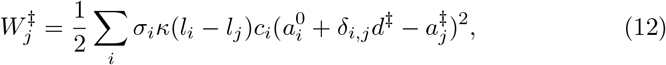

where *δ*_*i,j*_ is 0 for *i* ≠ *j* and 1 for *i* = *j*. We could thus calculate the mechanical work difference,

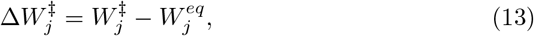

required to change the conformation of a given protein *j* by 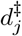, and attain the associated local mechanical equilibrium.

We will now proceed to describe how changes in mechanical work alter the rates of molecular transitions, namely binding, mechanical steps, or unbinding of proteins. For reactions that are not load-dependent (myosin and filamin attachment to actin), we assumed rates that are independent of mechanical load (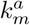 and 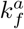, respectively). For all other, load-dependent reactions we assumed that a conformational change 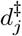 of the myosin or filamin protein *j* would have to occur to reach the transition state and initiate the reaction. For a given load-dependent transition, we assumed that 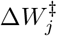 affects the transition rate as

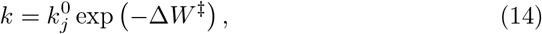

where 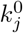 is the unloaded transition rate for a given reaction. Note that the mechanical work is given in units of *k*_*B*_*T*, so that the typical *W*/*k*_*B*_*T* normalization in the exponent is omitted [9].

For myosin binding sites, we used the same reaction scheme as in our previous work [9], consisting of a unidirectional cycle of myosin binding, main power stroke, and minor power stroke leading to detachment. Myosin attachment occurred with rate

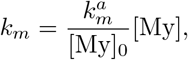

where 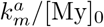 captures both the effective attachment rate of myosin on the coverslip to the actin filament, as well as the influence of soluble myosin concentration ([My]) on this rate. Upon binding to actin, the myosin at a given binding site *j* was assigned an initial strain from a normal distribution with a standard deviation *w*_*m*_. The main power stroke occurred with an unloaded rate 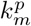, and always had a step length 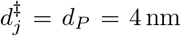. The minor power stroke occurred with an unloaded rate 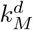, and always had a step length 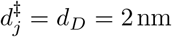. The effective spring constant of myosin was *c*_*m*_.

Filamin attachment occurred with a rate 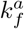 at random positions 0≤*l*_*j*_≤*L* along the actin filament, and without pre-existing strain 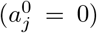. Filamin detachment occurred with an unloaded rate 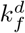. The distance from the current position to the transition state required for detachment was calculated based on the current configuration,

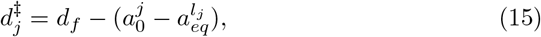

where *d*_*f*_ is the total distance from relaxed filamin to a stretched state required for detachment.

#### 3.2 Numerical evaluation and extraction of motion traces

To numerically simulate the mechanochemical interactions of actin, myosin, and filamin, we extended our previously developed algorithm for the interaction of myosin with actin [9, 5]. First, in the reaction scheme that we used, for each individual myosin binding site and filamin protein, always only one reaction step was possible as the next reaction. Hence, a single rate for a reaction to occur for any of the *j* simulated elements could be calculated, under the assumption of local mechanical equilibria. Second, these rates were used in a Gillespie algorithm step to determine the waiting time until the next transition occurs as well as the element for which the next transition occurs [9]. The simulation time was incremented by the waiting time, and the variables associated with the altered element (chemical state, *α*_*j*_, 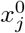) were updated. Third, the macroscopic position of the actin filament was extracted by calculating a global *a* under the assumption *κ* = 1, so that all proteins bound to the actin filament contribute equally, independent of their position. In this manner, a single position of the actin filament could be extracted, but no change to the microscopic state of the mechanochemical system was effected. It is clear that some inaccuracies will result from this *ad hoc* approach to extracting the actin position. However, these should be well below ≈ 100 nm. Our goal was a comparison with experimental tracking data, and inaccuracies of 100 nm can easily result from recording noise, tracking inaccuracies, or fluctuations in the actin sliding progress in the experiment (see below). Based on these considerations, we assign microscopic accuracy only to simulations assuming local mechanical equilibria. The assumption of global equilibrium (*κ* = 1) served purely as an *ad hoc* approach to obtain a scalar actin displacement value that can be compared to our experimental observations. Note, again, that the calculations with *κ* =1 did not feed back on the microscopic mechanochemistry in the simulation.

To further make the simulation results comparable with experimental data, which were recorded with a fixed sampling interval Δ*t* = 0.33 s, the simulation data were resampled at the same time interval Δ*t* [9]. *V*_*f*2*f*_ values were calculated from the differences in *a* at consecutive sample times, divided by Δ*t*. To account for inevitable noise in filament sliding progress present in the experiment, we added a random number drawn from a normal distribution with standard deviation 0.1 *μ*m/s to each *V*_*f*2*f*_ value (appropriate value for Δ*t* = 0.33 s) [5].

#### 3.3 Choice of model parameters

The model parameters used in the detailed model (SI Tab. 1) were taken from existing literature were possible, and adjusted to our experimental data where no previous information existed. The model parameters for interactions of only actin and myosin in the absence of filamin were taken from our previous work [4]. Assuming initially *L*_*c*_ → ∞, the model parameters 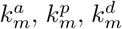, *c*_*m*_, and *w*_*m*_ were adjusted to the experimental results for short actin filaments (*L* < 0.8 *μm*) at the highest myosin concentration ([My] = 0.166 mg/ml), see SI Fig. 1. This step of parameter adjustment was based on an analysis of the influence of the different parameters from our previous work [5]. Note that 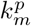 and 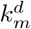 were both measured in single myosin experiments previously, yielding values of ≈ 200 s^−1^ and ≈ 25 s^−1^ at 23°C, respectively, which are close to the values we obtained from our adjustment [10]. The myosin step lengths were also taken directly from single molecule experiments [11]. Using these parameters, simulations with different *L*_*c*_ < ∞ were carried out, and *L*_*c*_ = 0.3 *μ*m was chosen for the best agreement with the experimental data in terms of the sliding velocity plateau for *L* ≫ *L*_*c*_ (SI Fig. 1B). This value of 0.3 *μ*m is physically justifiable, as it is above a lower-bound distance of ≈ 130 nm, which is where the compound stiffness of actin-attached myosin can first exceeds actin longitudinal stiffness (see Discussion in main manuscript).

The mechanical parameters of single filamin crosslinks 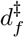 and *c*_*f*_, were chosen within the range of published results [12]. A ratio *c*_*f*_/*c*_*m*_ ≈ 300 was applied, to reflect the axial stiffnesses of 880 ± 551 pN/nm of filamin [12] and ≈ 2.9 pN/nm of the myosin S1 region [13]. Note the stiffness values for the taut chain configuration are used for both proteins, to reflect that they take effect in our simulations only once taut. The only left-over free parameters, 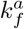 and 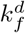 were adjusted to *f*_*mot*_ from our experimental data (Fig. 5D), and produced *τ*_*AC*_ values that were in good agreement with our experimental observations (Fig. 5E).

**Tab. 1.** Parameters of the detailed model.

| Parameter | Symbol | Value | Source |
| --- | --- | --- | --- |
| **Actin-myosin interaction** |  |  |  |
| Attachment rate | $k_m^a$ | $163.7 \text{ s}^{-1} (\text{mg/ml})^{-1}$ | Our work, compare Hilbert *et al.*[4] |
| Power stroke rate | $k_m^p$ | $256.5 \text{ s}^{-1}$ | Our work, compare Veigel *et al.*[10] |
| Detachment rate | $k_m^d$ | $15.2 \text{ s}^{-1}$ | Our work, compare Veigel *et al.*[10] |
| Myosin effective spring constant | $c_m$ | $3.5 \text{ nm}^{-2}$ | Hilbert *et al.*[4] |
| Myosin attachment range | $w_m$ | $4.5 \text{ nm}$ | Hilbert *et al.*[4] |
| Main power stroke length | $d_P$ | $4 \text{ nm}$ | Capitanio *et al.*[11] |
| Minor power stroke length | $d_D$ | $2 \text{ nm}$ | Capitanio *et al.*[11] |
| **Longitudinal decay of mechanical coupling strength** |  |  |  |
| Coupling strength decay length | $L_c$ | $0.3 \mu\text{m}$ | Our work |
| **Actin-filamin interaction** |  |  |  |
| Filamin attachment rate | $k_f^a$ | $1 \text{ s}^{-1}$ | Our work |
| Filamin detachment rate (unloaded) | $k_f^d$ | $100 \text{ s}^{-1}$ | Our work |
| Filamin effective spring constant (when taut) | $c_f$ | $1000 \text{ nm}^{-2}$ | Ferrer *et al.*[12] |
| Distance to detachment transition state (when taut) | $d_f$ | $0.2 \text{ nm}$ | Ferrer *et al.*[12] |

### 4 Spin chain model

#### 4.1 Development of the spin chain model

A linear chain of spins *s*_*n*_ (*n* = 1, 2,…, *N*) represents the major myosin binding sites distributed along the actin filament. *N* = *L*/*b* was used in the same manner as for the detailed model to chose the number of spins corresponding to a given actin length *L*. Each spin can take the value +1 or −1, representing a myosin binding site associated with the active or the arrested myosin group state, respectively.

The spin value changes stochastically, with a switch rate depending on the value of the spin itself, *s*_*n*_, as well as its nearest neighbors, *s*_*n*−1_ and *s*_*n*+1_,

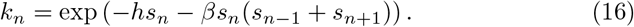

Here, *h* is a global field, and *h* > 0 indicates a tendency towards *s*_*n*_ = +1 imparted by this field. *β* describes the coupling strength between the nearest neighbors, with *β* > 1 favoring nearest neighbors with the same spin value. For the first (*n* = 1) and the last (*n* = *N*) spin in the chain, the influence of the nearest neighbors is calculated differently. Values for *s*_0_ and *s*_*N*+1_ are required, but not contained in the spin chain. Instead

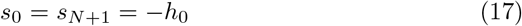

is used. Thus, *h*_0_ > 0 would introduce a bias towards the *s*_*n*_ = −1 state at the ends of the spin chain.

#### 4.2 Exact calculation of the motile fraction

In the in vitro motility assay, *f*_*mot*_ is determined as the fraction of time during which actin motion faster than a given threshold velocity is observed. This measurement is mirrored here by calculating the probability that all *s*_*n*_ = +1,

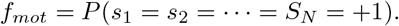

This assumption is based on the mechanistically detailed simulation, where local arrest of the myosin kinetics leads to global arrest of actin sliding.

The exact value of *f*_*mot*_ was calculated for an ensemble of equilibrated spin chains. The probability to encounter a given state **s** = (*s*_1_, *s*_2_,…, *s*_*N*_) can be calculated based on the potential energy associated with s,

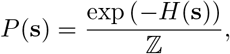

where

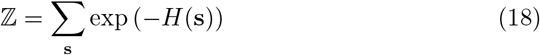

is the partition function, found by summation over all possible configurations of **s**.

The potential energy of a state **s** can be calculated based on the kinetic rates specified above. Specifically, for an individual spin *s*_*i*_, the contribution to the potential energy is,

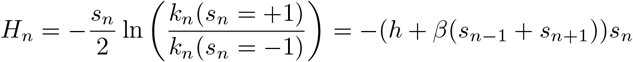

By summation across all spins, the overall potential energy of a given state **s** can be calculated,

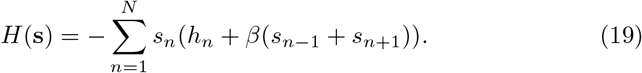

It is now possible to write down

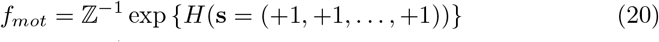

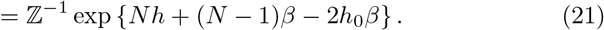

For efficient evaluation, the partition function can be written in the form of scalar products

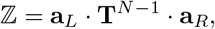

Where

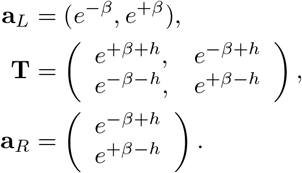

#### 4.3 Introduction of crosslinkers

To introduce crosslinkers, which can bias the spins into the *s*_*n*_ = −1 state, we included for each spin an additional binary variable *c*_*n*_ ∈ {0, 1}. *c*_*n*_ = 1 referred to the presence of a crosslinker at the position of the respective spin, and affected the spin switching rate as

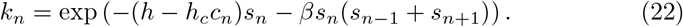

Here, *h*_*c*_ > 0 quantifies the bias towards the *s*_*n*_ = −1 state that is exerted by the presence of a crosslinker. The rate for a state change of the crosslinker variable *c*_*n*_ is

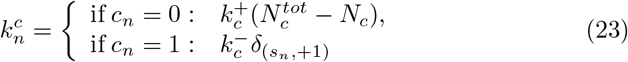

Here, 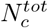 refers to the total number of crosslinkers available, and *N*_*c*_ to the number of crosslinkers currently bound. 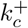 and 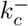 are the crosslinker binding and unbinding rates, respectively. 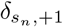 is 0 if *s*_*n*_ = −1 and 1 if *s*_*n*_ = +1, to reflect the detachment of crosslinkers by active myosin seen in the detailed model.

#### 4.4 Stochastic simulation of spin chain model

To produce example plots of spin chain kinetics, extract autocorrelation times, and to evaluate the spin chain model with crosslinkers, the rate expression were used to carry out Gillespie simulations [14]. Time courses of average activity were extracted by sampling

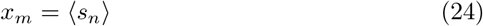

at different time points *m* separated by Δ*t* = 0.1. *f*_*mot*_ was then calculated as the fraction of all *x*_*m*_ that had a value of 1, corresponding to full activity. To calculate the autocorrelation time, to each *x*_*m*_ value a random variable sampled from a uniform distribution between 0 and 0.05 was added, then the autocorrelation time was extracted.

#### 4.5 Choice of model parameters

The fitting of the model parameters that specified the spin chain model without inhibiting crosslinkers (*β*, *h*, and *h*_0_) was executed based on a minimization of the root of the sum of the squared error (*ε*) between *f*_*mot*_ from model and experiment. First, *h*_0_ = 0.1 was assigned and a range of *β* ∈ [0, 10] and *h* ∈ [0, 0.5] was scanned by brute force to choose the (*β, h*) combination that minimized *ε*. Second, *h* was released, and the Nelder-Mead simplex (direct search) method was employed to further minimize *ε* by adjusting (*β*, *h*, *h*_0_), starting from the results of the brute force search. For the addition of crosslinkers, the parameters *h_c_* = 1.5, 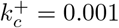, and 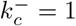 were assigned to qualitatively reproduce experimental *f*_*mot*_ and *τ*_*AC*_ curves.

